# A *Vibrio cholerae* Type IV restriction system targets glucosylated 5-hydroxyl methyl cytosine to protect against phage infection

**DOI:** 10.1101/2024.04.05.588314

**Authors:** Jasper B. Gomez, Christopher M. Waters

## Abstract

A major challenge faced by *Vibrio cholerae* is constant predation by bacteriophage (phage) in aquatic reservoirs and during infection of human hosts. To overcome phage predation, *V. cholerae* has evolved a myriad of phage defense systems. Although several novel defense systems have been discovered, we hypothesized more were encoded in *V. cholerae* given the relative paucity of phage that have been isolated which infect this species. Using a *V. cholerae* genomic library, we identified a Type IV restriction system consisting of two genes within a 16kB region of the *Vibrio* pathogenicity island-2 that we name TgvA and TgvB (**<u>T</u>**ype I-embedded **<u>g</u>**mrSD-like system of **<u>V</u>**PI-2). We show that both TgvA and TgvB are required for defense against T2, T4, and T6 by targeting glucosylated 5-hydroxymethylcytosine (5hmC). T2 or T4 phages that lose the glucose modification are resistant to TgvAB defense but exhibit a significant evolutionary tradeoff becoming susceptible to other Type IV restriction systems that target unglucosylated 5hmC. We show that additional phage defense genes are encoded in VPI-2 that protect against other phage like T3, secΦ18, secΦ27 and λ. Our study uncovers a novel Type IV restriction system in *V. cholerae*, increasing our understanding of the evolution and ecology of *V. cholerae* while highlighting the evolutionary interplay between restriction systems and phage genome modification.

**Abstract Importance:** Bacteria are constantly being predated by bacteriophage (phage). To counteract this predation, bacteria have evolved a myriad of defense systems. Some of these systems specifically digest infecting phage by recognizing unique base modifications present on the phage DNA. Here, we discover a Type IV restriction system encoded in *V. cholerae* that we name TgvAB and demonstrate it recognizes and restricts phage that have 5-hydroxymethylcytosine glucosylated DNA. Moreover, the evolution of resistance to TgvAB render phage susceptible to other Type IV restriction systems, demonstrating a significant evolutionary tradeoff. These results enhance our understanding of the evolution of *V. cholerae* and more broadly how bacteria evade phage predation.

## Introduction

*Vibrio cholerae,* the causative agent of cholera, is an acute diarrheal disease that has greatly impacted human history. Today, it is estimated that 1.3 to 4 million *V. cholerae* infections occur annually worldwide leading to around 21,000 to 143,000 deaths(1, 2). A major evolutionary selection *V. cholerae* must overcome is predation by bacteriophages (phage) in both its aquatic environment and pathogenic lifestyle(3). *V. cholerae* epidemics are intricately associated with phage infections as phage blooms curtail cholera outbreaks(4). Three dominant lytic phages, ICP1, ICP2, and ICP3 have been identified to eradicate *V. cholerae* both *in vivo* and during environmental persistence(3). However, given the ubiquitous nature of *V. cholerae* in environmental reservoirs, it is curious that a more diverse population of phage targeting this bacterium has not been discovered, suggesting that *V. cholerae* is adept at defending against phage infection.

To overcome phage predation, *V. cholerae* has evolved a myriad of phage defense systems(5–8). Interestingly, these phage defense systems are intricately linked to genomic regions or prophages acquired by horizontal gene transfer including the *Vibrio* pathogenicity islands (VPI-1 and VPI-2)(9, 10) and *Vibrio* seventh pandemic island (VSP-1 and VSP-2)(11, 12). The VSP-1 and VSP-2 islands are only found in the current 7^th^ pandemic El Tor biotype strains, and their association with phage defense supports that these islands allow for long-term persistence and potentially the emergence of this pandemic(13–15). VSP-1 encodes a recently identified phage defense system, *avcID*(6). AvcI is a noncoding RNA antitoxin that inhibits the cytidine deaminase activity of AvcD(6). Upon the inhibition of host gene transcription, AvcI is rapidly degraded, liberating AvcD to deplete free deoxycytidine nucleotides within the cell, decreasing phage replication(16). VSP-I also encodes *dncV, capV, vc0180* and *vc181,* a **<u>c</u>**yclic-oligonucleotide-**<u>b</u>**ased **<u>a</u>**nti-phage **<u>s</u>**ignaling **<u>s</u>**ystem (CBASS) phage defense system, that limits phage infection to the surrounding population through abortive infection(8, 17, 18). VSP-2 encodes the genes *ddmABC,* which are DNA condensins that have been shown to limit both phage and plasmid replication(7, 12). A partner system, *ddmDE,* that also limits plasmid acquisition and potentially phage infection is encoded on VPI-2(7). *V. cholerae* also encodes a phage-inducible chromosomal-like element (PLE) that inhibits replication of ICP1 by confiscating its replication machinery(19–21)

Given these diverse defense systems, it is interesting that restriction/modification systems (R-M), which have been appreciated for decades as mediating phage defense(22), have not been described in *V. cholerae*. These enzymes are grouped into four classes. Canonical restriction enzymes that belong to Classes I-III, which are widely used in molecular biology, digest unmodified DNA at a specific sequence(23, 24). The DNA of the host is protected by a cognate methylase that blocks the activity of the restriction enzyme whereas unprotected phage DNA is rapidly degraded upon infection(23, 24). In response to these defense systems, phage have evolved mechanisms to modify the bases of their genomes(25–29). For example, T-even coliphage (T2, T4, and T6) all specifically methylate their cytosines at the 5’ position to generate 5-hydrocymethylcytosine (5hmC), which blocks the activity of many R-M systems(30).

Such modification selected for the Type IV class of restriction enzymes (24, 27, 31–33). Rather than targeting unmodified DNA, these enzymes specifically digest modified DNA associated with phage. For example, the McrA and McrBC restriction systems of *E. coli* digests genomes with either 5-methylcytosine (5mC) or the 5hmC modification(31, 34, 35). In response to restriction enzymes like McrA and McrBC, phage evolved additional base modifications to prevent digestion such as gluocosylation, which adds either a- or b-glucose to 5hmC. However, other Type IV restriction enzymes, as demonstrated by GmrSD, evolved to specifically digest glucosylated 5hmC(36–39). Therefore, bacteria and phage have complex evolutionary interplay between phage defense mediated by restriction enzymes and phage counter defense strategies that further alter the phage genome.

Given the relative paucity of phages that have been identified to infect *V. cholerae,* we hypothesized that this bacterium encodes previously undescribed phage defense systems. To identify such systems, we implemented a unbiased forward genetic screen of a *V. cholerae* genomic cosmid library for genes that protected *Escherichia coli* from phage defense. Using this approach, we identified the genes *vc1767* and *vc1766*, renamed herein as *tgvAB* for **T**ype I-embedded ***g****mrSD*-like system of **V**PI-2, provided broad protection against T-even phage. We find that TgvAB specifically targets phage DNA with glucosylated 5-hydrocymethylcytosine (5hmC) bases. Phages can evolve to overcome TgvAB by null mutations in their *agt* and/or *bgt* genes encoding the α- or b-glucosyltransferase that glucosylates 5hmCs. However, phage that evolve resistance to TgvAB exhibit a significant tradeoff that leaves them vulnerable to restriction by the Type IV Restriction System, McrA and McrBC, which targets unglucosylated 5hmC. Our study adds restriction enzymes to the phage defense repertoire of *V. cholerae* and demonstrates the inherent evolutionary interplay between phage genome modification and restriction defense.

## Results

### A region within VPI-2 provides defense against phage

To identify novel phage defense systems in *V. cholerae,* we screened a *V. cholerae* genomic library encoding random 25 kB fragments from *V. cholerae’s* genome inserted into the cosmid pLAFR for protection against phage infection in *E. coli*. We screened for phage defense activity in *E. coli* as we hypothesized the dominant vibriophages, ICP1, ICP2, and ICP3, may be inherently resistant to such systems or have evolved counter defense mechanisms, which contributes to their dominance. Additionally, we have shown that *V. cholerae* phage defense systems such as CBASS and AvcID can be expressed in *E. coli* and confer protection against well studied coliphage(6, 17). We, therefore, screened ∼350 individual clones encoding the *V. cholerae* genomic library for resistance to T2 phage infection using an efficiency of plaquing (EOP) assay. We chose T2 as our initial phage to screen as it demonstrates sensitivity to *V. cholerae* CBASS in *E. coli*(17). One cosmid, which we named pJBG007, conferred complete protection against T2 infection as compared to the sensitive pLARF cosmid control (Fig. 1A). Additionally, infection of T2 at a multiplicity of infection (MOI) of 0.01 in liquid cultures of *E. coli* indicated that pJBG007 provided significant protection compared with pLAFR (Fig. 1B).

**Fig. 1.**
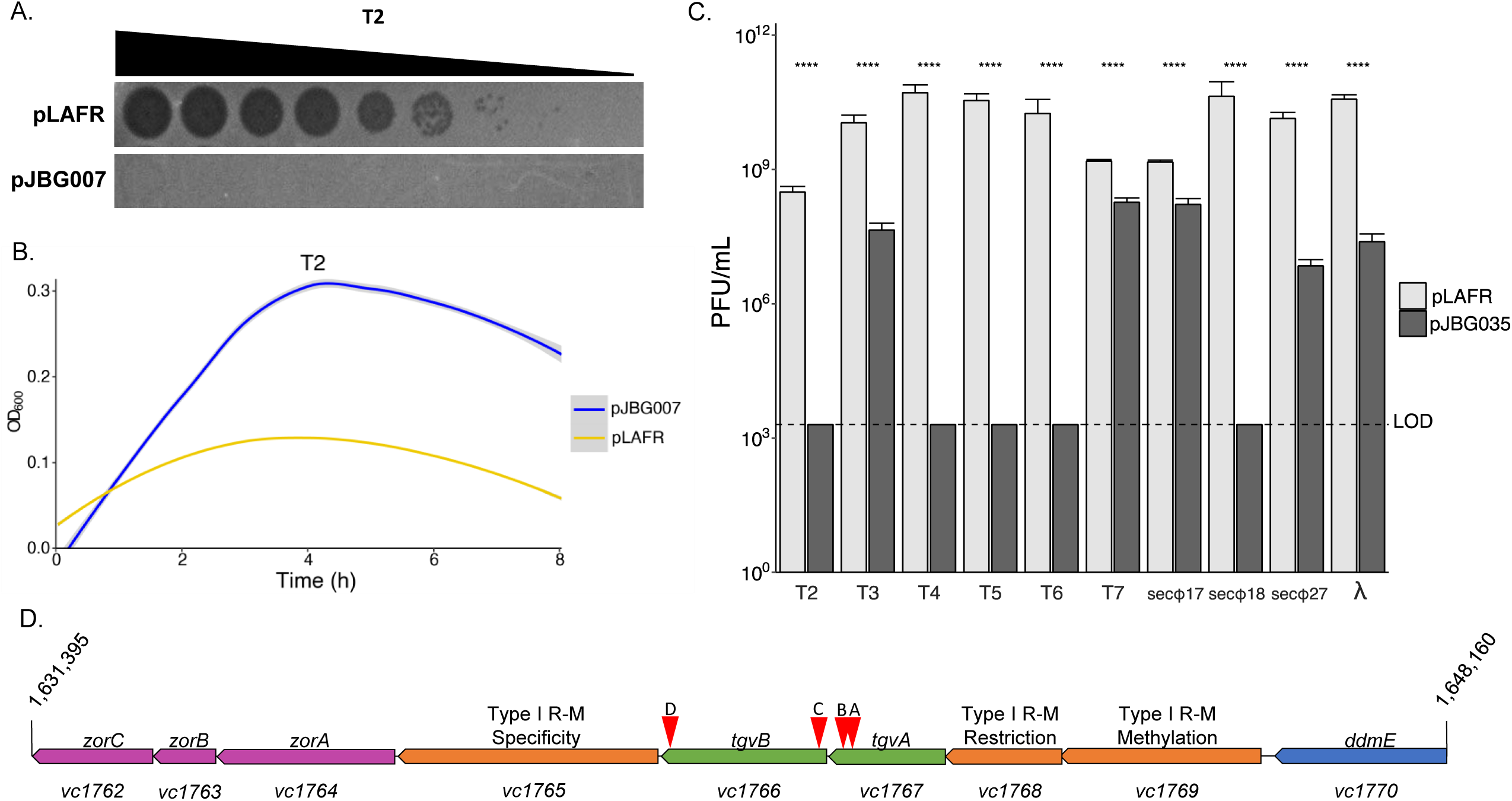
*V. cholerae* El Tor contains phage defense systems on VPI-2. **A.** 10-fold serial dilution plaque assay of T2 phage spotted on pJBG007 a cosmid containing *vc1770-vc1762* compared to pLAFR (empty vector). **B.** *E. coli* grown with either empty vector (yellow line) or pJBG007 (blue line) infected at time 0 with T2 phage at an MOI of 0.01. The mean and standard error of 4 biological replicates each with 3 technical replicates are presented. **C.** Plaque forming units (PFU/mL) for the indicated phage were spotted on *E. coli* containing empty vector (light gray) or pJBG007 (dark gray). The dashed line denotes the limit of detection (LOD). A Welch’s t-test was used to compare the mean (7 biological replicates each with 3 technical replicates) PFU count for each phage between empty vector and pJBG007. **D.** A schematic of VPI-2 region within pJBG007, highlighting transposon insertions A, B, C, and D for Fig. 2 is shown.

To test the breadth of pJBG007 protection against other phage, we quantified infection of 9 additional coliphage against *E. coli* containing pJBG007. Quantification of plaque forming units (PFU) indicated pJBG007 provided complete protection compared to pLAFR against T2, T4, T5, T6, and secΦ18, with no observable plaques at any dilution analyzed. Moreover, moderate but statistically significant protection against T3, secΦ18, secΦ27 and λ was also observed (Fig. 1C). These results indicated that the pJBG007 encodes genes that provide broad and significant protection against coliphage infection.

### pJBG007 encodes a novel *V. cholerae* phage defense cluster

Sequencing of the genomic DNA present in pJBG007 revealed that a 16,765 basepair fragment encoded in the cosmid was part of the Vibrio Pathogenicity Island-2 (VPI-2) (Fig. 1D). VPI-2 encodes the *ddmDE* biological defense system(7), but only *ddmE* was present on pJBG007. Previous results have shown that both *ddmD* and *ddmE* are required for function of this system, indicating that *ddmE* likely was not contributing to the defense we observed(7). In addition to *ddmE*, this fragment encoded an annotated Type-I R-M, and a truncated newly described phage defense system known as Zorya (Fig. 1D), neither of which have been studied in *V. cholerae*. Based on sequence homology, these genes would be considered a Type-I Zorya system. Prior studies suggest Type-I Zorya systems encode four genes, *zorABCD*, of which only *zorABC* are present in pJBG007(40). Therefore, it is currently unclear if *vc1764-vc1762* encodes a functional Zorya defense module. Phage defense systems cluster in defense islands(40), and therefore this fragment of DNA is a relatively unstudied defense island of *V. cholerae.* Recognizing that either the R-M system or Zorya system could be causing protection responsible for phage protection, we sought to identify the genes necessary for protection against T2.

### *vc1766 and vc1767* are necessary and sufficient to inhibit infection by T-even phages

To determine which genes were responsible for protection against T2 infection, we mutagenized pJBG007 with a transposon and tested ∼50 transposon mutants for loss of defense against T2 infection. We identified 4 transposon mutants in pJBG007 that lost all defense to T2 (Fig. 2A). These 4 transposon mutants also failed to protect *E. coli* infected in a liquid culture from T2 at an MOI of 0.01 (Fig. 2B). Sequencing of these transposon mutants identified two independent insertions in *vc1767* and two independent insertions in *vc1766* (Fig. 1D).

*vc1767* and *vc1766* are in a putative operon between genes *vc1769-vc1765* that was annotated as a Type-I R-M system (Fig. 1D). Basic Local Alignment Search Tool (BLAST) analysis indicated that *vc1767* and *vc1766* are both part of the COG1479 protein family, which is a conserved family of Type IV restriction DNase/DNA nickases specific for phosphorothioated or glycosylated phage DNA. This family is represented by GmrSD, which targets glucosylated 5hmC bases and DndB/SspE, which targets phosphorothioated bases(28, 41). *vc1767* and *vc1766* both encode predicted DUF262 which sense modified DNA and HNH nuclease domains. Given this analysis and the results described below, we renamed *vc1767* and *vc1766* to **<u>T</u>**ype I-embedded **g***mrSD*-like system of ***V***PI-2, *tgvA* and *tvgB*, respectively.

**Fig. 2.**
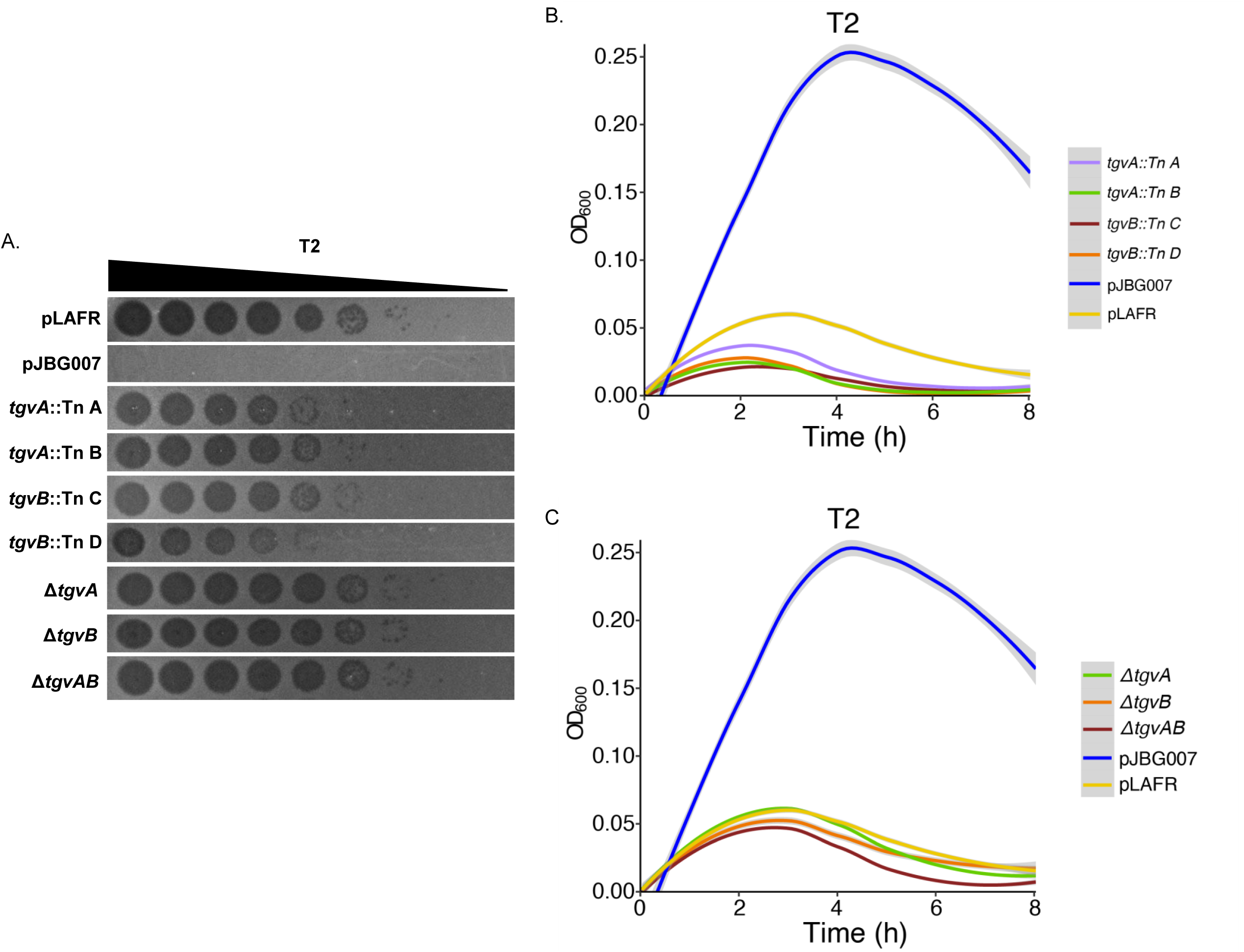
Disruption of *tgvA, tgvB,* and *tgvAB* causes loss of phage defense against T2. **A.** A 10-fold serial dilution plaque assay of T2 phage spotted on empty vector (pLAFR), pJBG007 (positive control), or the indicated mutants. **B.** Growth curves for the pLAFR, pJBG007 and the transposon mutants, or **C.** *tvgA* or *tvgB* deletion mutants infected at time 0 with T2 phage at an MOI of 0.01. The mean and standard error of 4 biological replicates each with 3 technical replicates are presented.

To test if the transposon insertions were polar on the surrounding genes, we generated D*tgvA,* D*tgvB,* or D*tgvAB* deletions in pJBG007 and measured T2 infection. Because the open reading frame (ORF) of *tgvA* overlaps the ORF of *tgvB* by 8 basepairs, we engineered the deletion of *tgvA* or *tgvB* such that the other ORF remained intact. Each of these mutants had a complete loss of protection (Fig. 2A). Similarly, T2 infection at an MOI of 0.01 in liquid cultures showed all three deletion mutants no longer protected against infection (Fig. 2C). Together, these results suggested that *tvgA* and *tvgB* were necessary for protection against T2 phage.

### TgvAB is sufficient for defense against T-even coliphages

To determine if *tvgA*, *tvgB*, or both genes were sufficient for phage defense, we created plasmids encoding *tgvA, tgvB,* and *tgvAB* expressed from their native promoter and screened for defense against T2 infection in *E. coli*. The p*tgvB* plasmid was made by amplifying a region 520 bp upstream of *tvgA* to downstream of the stop codon of *tvgB* using the D*tvgA* mutant as a DNA template such that *tvgB* would be expressed from its native promoter without *tvgA*. For controls, we used pJBG007, which provides robust defense, and the corresponding empty vector pEVS. Plasmids expressing the individual genes, p*tgvA* and p*tgvB,* exhibit plaquing equivalent to the empty vector, indicating these genes alone do not protect against infection. However, p*tgvAB* expressing both genes showed protection to T2 equivalent to pJBG007 (Fig. 3A). Analogous results were obtained in a liquid infection assay with T2 at an MOI of 0.01 indicating only p*tgvAB* provided protection from infection (Fig. 3B).

**Fig. 3.**
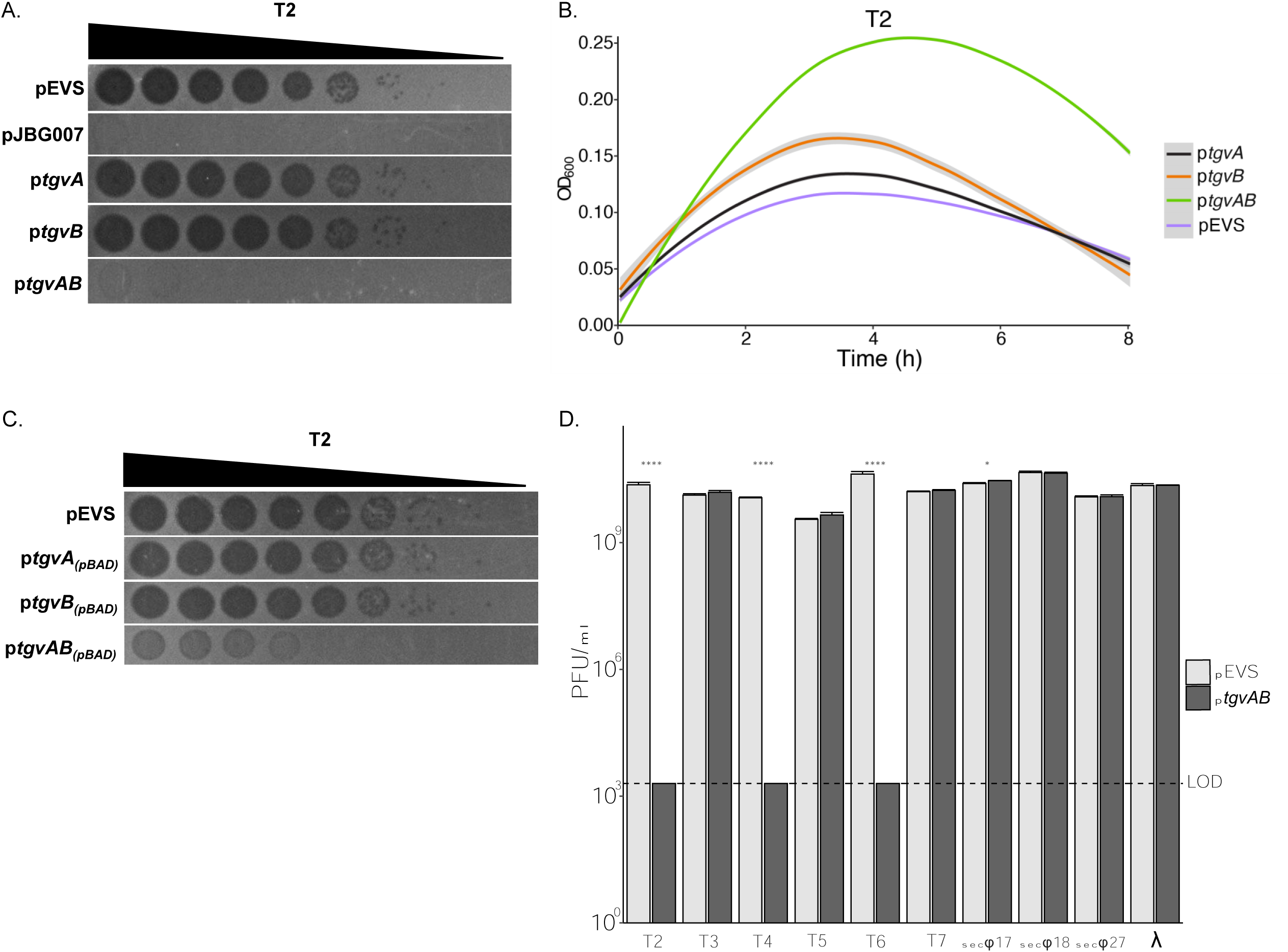
*tgvAB* confers protection against T2. **A.** A 10-fold serial dilution plaque assay of T2 phage spotted on pEVS (empty vector), pJBG007, p*tgvA*, p*tgvB* or p*tgvAB* expressing these genes from their native promoter is shown. **B.** Growth curves for the pEVS empty vector, p*tgvA*, p*tgvB*, and p*tgvAB*, infected at time 0 with T2 phage at an MOI of 0.01 are shown. The mean and standard error of 4 biological replicates each with 3 technical replicates are presented. **C.** T2 was spotted as a 10-fold serial dilution plaque assay on empty vector, p*tgvA****_(pBAD)_***, p*tgvB****_(pBAD)_***, and p*tgvAB****_(pBAD)_*** expressed using an arabinose inducible *P*_BAD_ promotor. **D.** The graph shows PFU/mL for T2, T4, T5, T6, T7, secΦ17, secΦ27 and λ infected on *E. coli* containing the empty vector (pEVS) or p*tgvAB*. The dashed line denotes the limit of detection (LOD). A Welch’s t-test was used to compare the mean (3 biological replicates each with 3 technical replicates) PFU count for each phage between empty vector and p*tgvAB*.

Due to the 8 base pair overlap between the *tgvA* and *tgvB* ORFs, we considered the possibility that expression of *tgvB* would be disrupted if *tgvA* was not present due to regulatory mechanisms like translational coupling. To account for such a possibility, we created overexpression plasmids encoding *tgvA, tgvB,* and *tgvAB* expressed from an arabinose inducible promoter and again screened for defense against T2 infection in *E. coli*. Plasmids overexpressing the individual genes, p*tgvA_(pBAD)_* and p*tgvB_(pBAD)_*, had plaquing equivalent to empty vector, again supporting that *tgvA* and *tgvB* expression alone was insufficient for protection. However, p*tgvAB_(pBAD)_*showed full protection, demonstrating that both genes are necessary and sufficient to provide phage defense whether they are expressed from their native promoters or exogenously induced (Fig. 3C).

Observing that p*tgvAB* was sufficient for protection against T2, we asked if *tgvAB* was responsible for the robust protection against T2, T4, T5, T6, and secΦ18 and moderate protection against T3, secΦ18, secΦ27 and λ that we observed for pJBG007 (Fig. 1C). We thus tested infection of p*tgvAB* and pEVS against all 10 coliphage and only observed robust protection against T2, T4 and T6 with p*tgvAB* (Fig. 3D). Based on these data, we conclude that *tgvAB* together is sufficient for protection against T-even coliphages, but other genes on pJBG007 are required for protection against T3, T5, secΦ18, secΦ27, and λ.

### T2 escapes TgvAB defense by mutations in α-glucosyltransferase (*agt*)

Our results thus far suggest that TgvAB is a Type IV restriction system that likely digests modified phage DNA. However, it was unclear which specific base modifications were targeted. To gain further understanding of *tgvAB* defense, we passaged T2 on *E. coli* containing p*tgvAB* in five independently evolved populations to isolate resistant mutations. These five populations were pooled together and monitored for susceptibility to p*tgvAB*. From this procedure, we isolated four T2 plaques from this pool that formed plaques on *E. coli* carrying p*tgvAB*.

In classical R-M systems, phage DNA passed in the presence of the modification enzyme can be modified such that it avoids restriction in subsequent infections. These modifications, however, are lost if phage is passaged in a host lacking the R-M system(42). To test whether these four individual T2 isolates escaped protection from *tgvAB* due to genome modifications mediated by *tgvAB*, we passaged all four individual T2 mutants on *E. coli* without *tvgAB* and isolated phage. We then infected *E. coli* containing p*tgvAB* or the empty vector (pEVS) and observed that all four T2 mutants still formed plaques in the presence of p*tgvAB* whereas WT T2 only plaqued in its absence (Fig. 4A). This result suggests that these phages had acquired resistant mutations.Similarly, all four T2 resistant mutants were insensitive to TvgAB in liquid culture (Extended Data Fig. 4A).

**Fig. 4.**
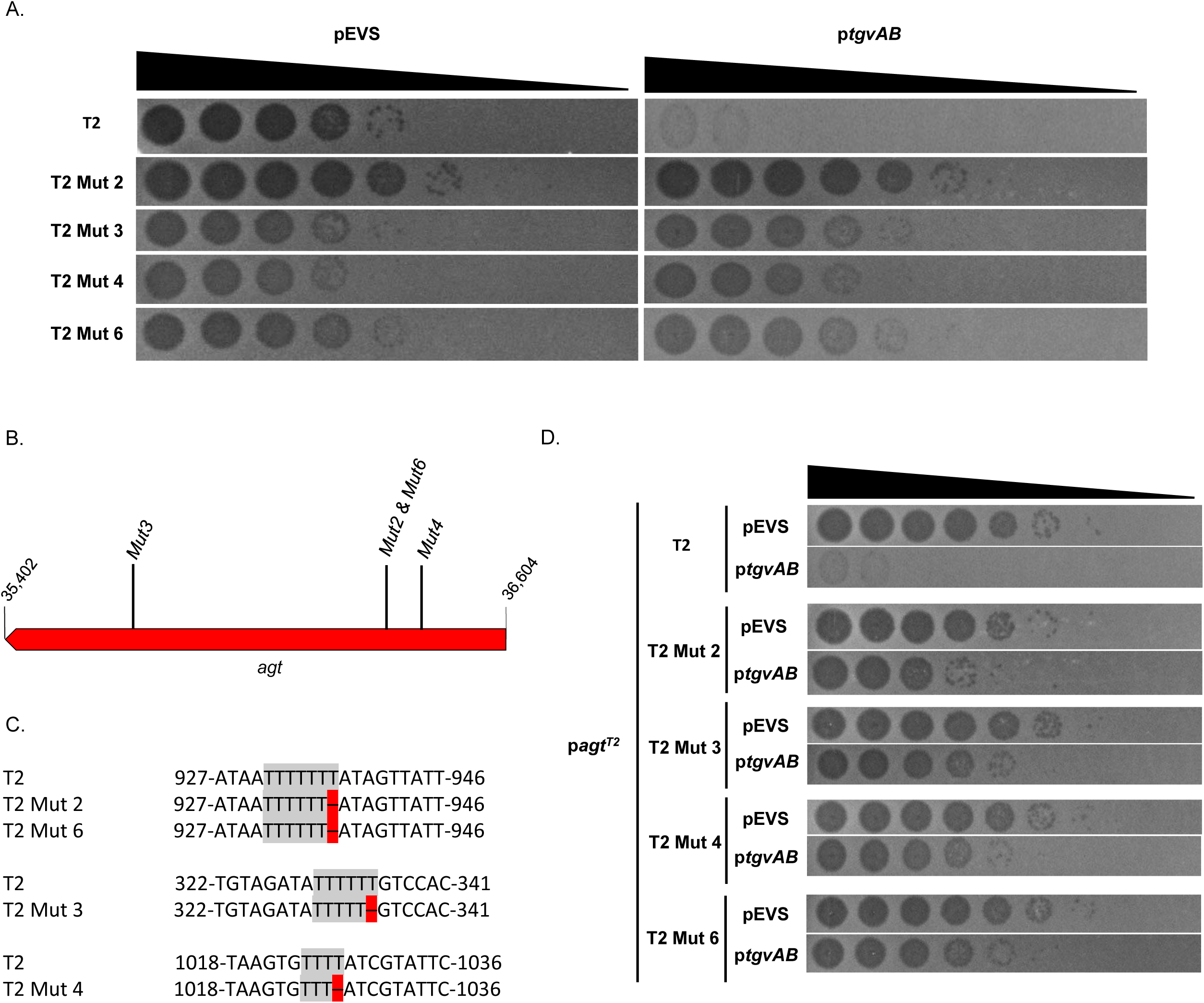
T2 escapes TgvAB defense by mutations in α-glucosyltransferase (*agt*). **A.** A plaquing assay of 10-fold serial dilution of WT T2 and evolved T2 mutants spotted on empty vector or a p*tgvAB* host. **B.** The Schematic of the *agt* gene in T2 indicates the position of the sequenced T2 mutations. **C.** The nucleotide sequence that is mutated and surrounding region in *agt* is compared to the sequence of *agt* from WT T2. Regions that are highlighted in red indicate a deletion of a T in that position. **D.** A plaquing assay of 10-fold serial dilutions of WT T2 and T2 Mutants passaged on host expressing p*agt*^T2^ with a *P_BAD_* promotor on empty vector or a p*tgvAB* host is shown.

Due to our analysis that predicted *tgvAB* belongs to the GmrSD/DndB/SspE family, all of which specifically digest phage DNA with unique modifications to 5-hydroxymethylctosine (5hmC)(26, 28, 43), we hypothesized that the T2 *tvgAB* resistant mutants had non-functional mutations in genes that were associated with unique modifications to 5hmC. We focused on the T2 gene *agt*, which encodes an α-glucosyltransferase that glucosylates 5hmC bases, due to prior studies showing the homologous GmrSD targets glucosylated phage DNA with these modifications and *agt* modifies 70% of the cytosines of T4 during an infection(28, 36). We sequenced the T2 *agt* gene for all four T2 mutants and a T2 control that was passaged in *E. coli* that did not have *tvgAB* and discovered each T2 resistant phage had a mutation in *agt* whereas the T2 control did not (Fig. 4B). Upon further analysis, all four mutants had a single nucleotide deletion of a thymine at three distinct poly-T tracts in the *agt* gene (Fig. 4C).

We then confirmed that all these mutants had deletion of a T using whole genome sequencing. The T2 Mut2 showed two additional mutations not found in T2 Mut6 indicating these were independent mutations even though they delete a T in the same poly-T track. Therefore, our sequencing analysis indicated that we isolated four independent *tvgAB* T2 resistant mutants that occurred in *agt,* and glucosylation of 5hmC of phage DNA is important for *tgvAB* defense.

To further explore the role of *agt*, we asked whether overexpression of T2 *agt* from a plasmid (p*agt*^T2^) in *E. coli* could restore glucosylation of 5hmC of the T2 resistant mutants, rendering them susceptible to *tgvAB*. All four T2 mutants and WT T2 were passaged in the *E. coli* strain containing p*agt*^T2^. As expected, WT T2 remained sensitive to p*tgvAB* as it encodes a functional *agt.* Passage of the T2 resistant phage in *E. coli* with p*agt*^T2^ increased susceptibility of these mutants to *tvgAB* by ∼10-100-fold (Fig. 4D). Taken together, these data suggest that TvgAB targets a-glucosylated 5hmC bases in T2 phage and loss of this modification renders T2 completely resistant to TvgAB.

### T4 Δ*agt*Δ*bgt* escapes TgvAB defense

Based on our previous results, we observed that *tgvAB* also confers protection against T4 and T6, each which have glucosylated 5hmC bases. We therefore explored the role of glucosylated 5hmC in *tgvAB* defense against T4. Indeed, liquid infection of T4 at a MOI of 0.01 in *E. coli* strains containing p*tgvA,* p*tgvB,* p*tgvAB*, and or the pEVS empty vector indicated that both *tvgA* and *tvgB* were required for phage defense, analogous to T2 (Fig. 5A). These results were confirmed using an EOP assay (Fig. 5B). We then asked whether T4 was able to escape *tgvAB* protection if it can no longer glucosylate 5hmC similar to the T2 TvgAB resistant mutants. To test this, we used a mutant T4 phage deleted for both *agt* and a second gene, *bgt,* that has β-glucosyltransferase activity on 5hmC (gift of Mike Laub). Prior studies have shown that both *agt* and *bgt* are required for glucosylation of 5hmC in T4 with *agt* a-glucosylating 70% of 5hmCs while *bgt* b-glucosylates 30% of these bases(36). Infection of *E. coli* containing p*tgvAB* or the empty vector (pEVS) with T4^Δ*agt*Δ*bgt*^ showed that this phage was unaffected by *tgvAB* (Fig. 5C). Restoration of glucosylation of 5hmC by passage of the T4^Δ*agt*Δ*bgt*^ mutant in *E. coli* strains overexpressing either T4 *agt* (p*agt*^T4^) or *bgt* (p*bgt*^T4^) restored partial susceptibility to *tgvAB* (Fig. 5C). Together, these data suggest that that either a- or b-glucosylation on 5hmC of T4 DNA is required for *tgvAB* recognition of T4, similar to our results for T2.

**Fig. 5.**
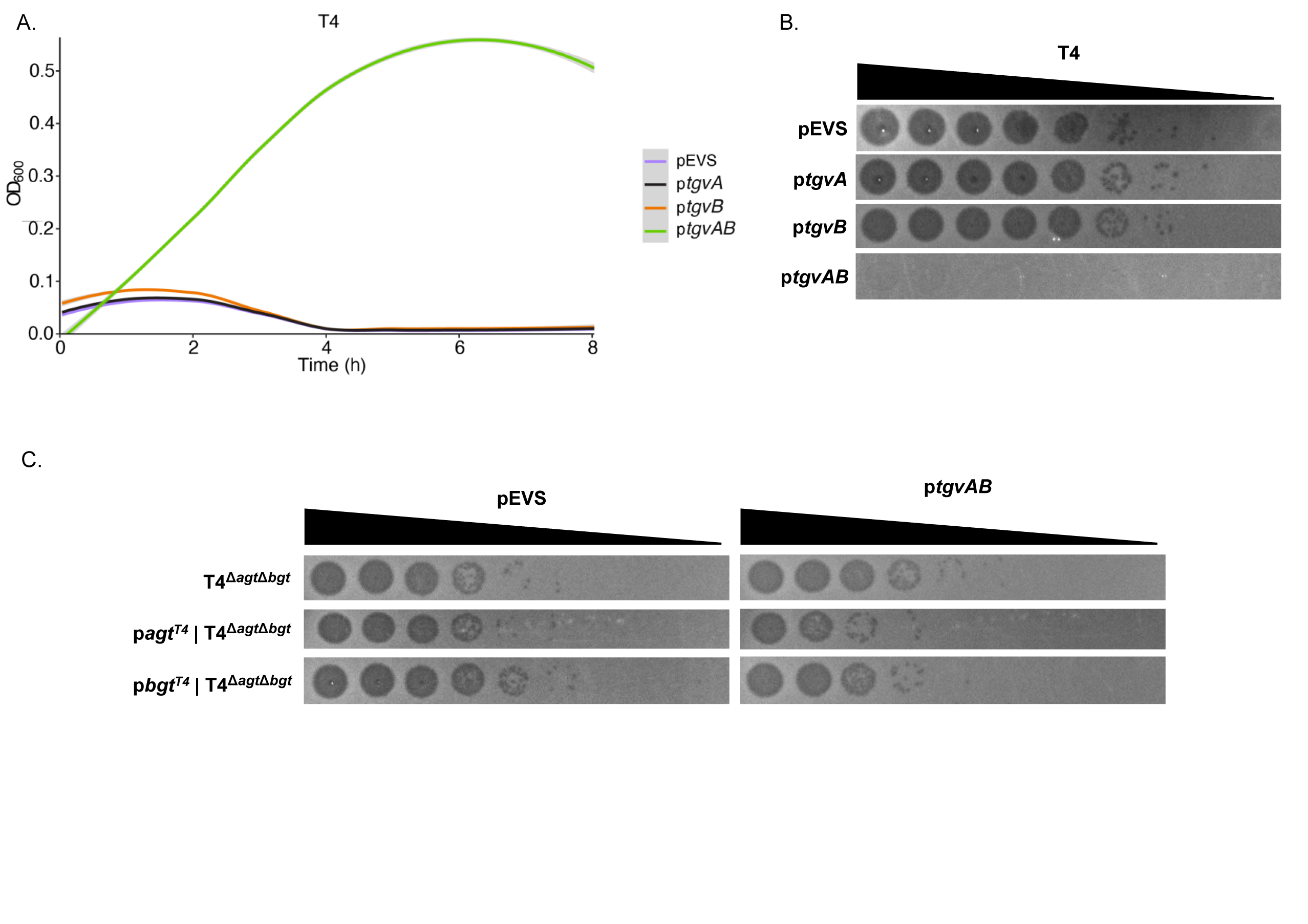
TgvAB protects against T4 infection and T4 Δ*agt*Δ*bgt* escapes TgvAB defense. **A.** *E. coli* encoding p*tgvA*, p*tgvB*, and p*tgvAB* was infected with T4 at an MOI of 0.01 at time 0. The mean and standard error of 4 biological replicates each with 3 technical replicates are presented. **B.** 10-fold serial dilutions of T4 were spotted on *E. coli* containing the empty vector (pEVS), p*tgvA*, p*tgvB*, and p*tgvAB*. **C.** 10-fold serial dilution of T4^Δ*agt*Δ*bgt*^ (top), T4^Δ*agt*Δ*bgt*^ were passed on host expressing p*agt*^T4^ (middle), or p*bgt*^T4^ (bottom) with a *P*_BAD_ promotor and spooted on *E. coli* containing the empty vector (pEVS) or p*tgvAB*.

### T2 and T4 mutants are susceptible to McrA and McrBC defense

Although the loss of *agt* and/or *bgt* may provide resistance to TgvAB defense, we hypothesized the unglucosylated 5hmC may now become sensitive to other Type IV restriction systems that digest unglucosylated 5hmC, such as the *mcrA* and *mcrBC* restriction systems encoded in *E. coli* strain M1655. We thus infected MG1655 with T2 and all four of the *tvgAB* resistance mutants. We observed that WT T2 showed substantial plaquing in MG1655; however, all the T2 resistant mutants exhibited little to no plaques (Fig. 6A). These results suggest MG1655 restricts the T2 *agt* mutants but is unable to restrict WT T2. This result also further demonstrates loss of glucosylation of 5hmC in our T2 resistance mutants.

**Fig. 6.**
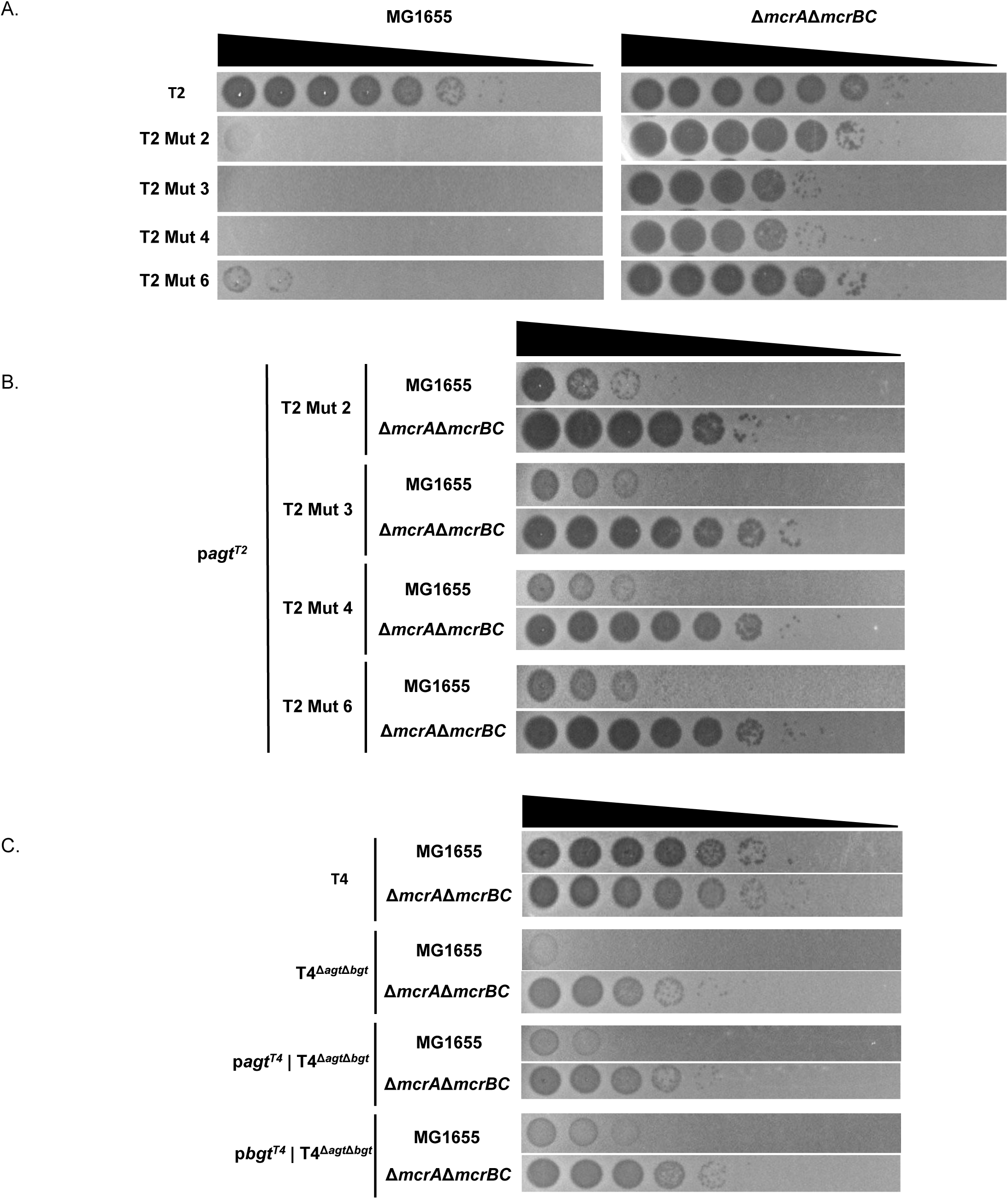
T2 and T4 mutants are susceptible to McrA and McrBC. **A.** 10-fold serial dilution of WT T2 and the four evolved T2 mutants were spotted on MG1655 WT or a MG1655 Δ*mcrA*Δ*mcrBC* host. **B.** The evolved T2 mutants were passaged on a host expressing p*agt*^T2^ then spotted as 10-fold dilutions onto MG1655 WT or a MG1655 Δ*mcrA*Δ*mcrBC* host. **C.** WT T4, T4^Δ*agt*Δ*bgt*^, T4^Δ*agt*Δ*bgt*^ were passed on host expressing p*agt*^T4^, or p*bgt*^T4^ with a *P*_BAD_ promotor then spotted as 10-fold dilutions on MG1655 WT and MG1655 Δ*mcrA*Δ*mcrBC* host.

We hypothesized that removal of *mcrA* and *mcrBC* would rescue the T2 resistant mutants’ infectivity. Indeed, infection of MG1655 Δ*mcrA*Δ*mcrBC* (gift of Mike Laub) with all T2 mutants and WT T2 showed robust plaquing (Fig. 6A). To test whether we could restore sensitivity of the T2 resistant mutants to *mcrA* and *mcrBC*, we infected MG1655 with T2 mutants passaged in the presence of p*agt*^T2^ and observed 100-1000-fold increased plaquing (Fig. 6B). These data further support that the T2 resistance *agt* mutants we isolated no longer glucosylate 5hmC and thus become sensitive to another Type IV restriction enzyme that targets unglucosylated 5hmC.

To determine if the above observations are specific to T2, we explored sensitivity of T4^Δ*agt*Δ*bgt*^ to *mcrA* and *mcrBC.* Infection of MG1655 with T4 and T4^Δ*agt*Δ*bgt*^ showed little to no plaques of T4^Δ*agt*Δ*bgt*^ whereas WT T4 exhibited substantial plaquing (Fig. 6C). However, infection of MG1655 with T4^Δ*agt*Δ*bgt*^ passaged on either p*agt*^T4^ or p*bgt*^T4^ showed 100-1000-fold increased plaquing (Fig. 6C). Thus, like T2, resistance to TvgAB via the loss of 5hmC glucosylation in T4 has an evolutionary tradeoff of increased sensitivity to other Type IV restriction systems like McrA/McrBC.

## Discussion

*V. cholerae* transitions between life in environmental reservoirs and infection of human hosts to cause cholera, but in each of these environments it has intricate interactions with phage(3). Although several phage defense systems have been discovered in this bacterium, our research demonstrates that there is still more to learn about how *V. cholerae* defends itself against phage infection.

VPI-2 is a pathogenicity island that is only present in toxigenic *V. cholerae* O1 and O139 serogroup isolates(9, 10). The acquisition of VPI-2 has been studied extensively; however, the biological function of most of the genes on VPI-2 remain unclear. Recently, a biological defense system called *ddmDE* was identified within VPI-2(7). Our unbiased forward genetic screen for novel *V. cholerae* phage defense systems identified a 16,765bp region of VPI-2 that protects *E. coli* from infection by a variety of phages (Fig. 1C). This region encodes only *ddmE* (without *ddmD*), an annotated Type I R-M System, and a newly described phage defense known as Zorya (Fig. 1D).

Prior studies have shown that there are two types of Zorya systems, type-I which encodes *zorAB* (*motAB* homologs), *zorC* (hypothetical protein) and *zorD* (large protein with helicase domain) and type-II which encodes *zorAB,* and *zorE* (gene encoding HNH-endonuclease)(40). We hypothesize that pJBG007 encodes a truncated type-I Zorya as it only encodes *zorAB,* and a hypothetical protein that shows no homology to *zorE*. Our results show that the previously annotated Type I Restriction system actually encodes two distinct restriction systems that potentially mediate phage defense. We demonstrate that a Type IV restriction system, which we name TgvAB, embedded within the annotated Type I Restriction System confers substantial protection against T2, T4 and T6 (Fig. 2B). We show that this protection is mediated through targeting of glucosylation of 5hmC modified bases, and loss of this gluosylation renders T2 and T4 completely resistant to TgvAB (Fig. 7).

**Fig. 7.**
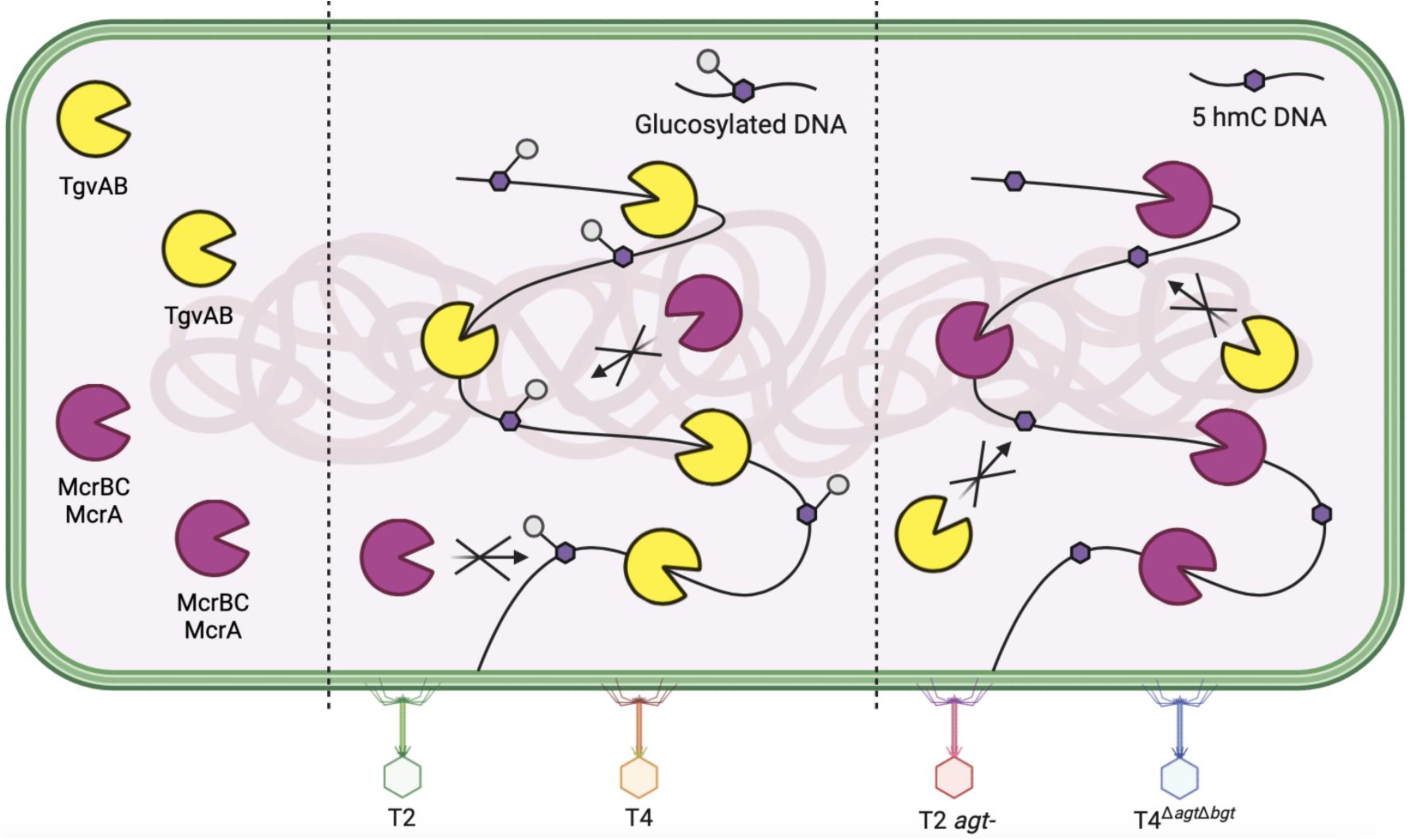
Model for TgvAB defense against phage and evolutionary tradeoffs of *agt* resistant mutants. In a cell expressing Type IV restriction systems like TgvAB, McrA and McrBC, when a phage adsorbs and injects its DNA, if the phage DNA is glucosylated at 5hmC bases, TgvAB recognizes this modification and restricts phage DNA to inhibit phage progeny (middle). If the phage DNA is injected and is not glucosylated, only having a 5hmC modification, it is resistant to TvgAB. However, McrA and McrBC recognize this mutant phage and restrict phage DNA inhibiting phage progeny (right).

Our analysis of this new phage defense region in VPI-2 encoded on cosmid pJBG007 demonstrated complete protection against T2, T4, T5, T6 and secΦ18 and significant protection to several other coli phage. However, as TgvAB alone only defended against T2, T4, and T6, the remaining genes in the Type I Restriction Modification and/or Zorya systems likely mediate defense to other phage, a possibility which we are exploring. This cluster of defense systems within VPI-2 support prior studies that have found phage defense systems frequently cluster together in phage defense islands(40). Furthermore, the identification of this phage defense cluster supports the notion that acquisition of VPI-2 contributed to long-term environmental persistence of *V. cholerae*(44).

Our results indicate *E. coli* carrying only *tgvAB* showed substantial protection against T-even phage infection by measuring both plaque forming units (PFU) and infection in a liquid culture. However, neither *tgvA* nor *tgvB* alone, when expressed from either their native promoter or induced with the pBAD promoter, exhibited significant phage protection. BLAST analysis of these genes showed that TgvA and TgvB belong to a conserved family of Type IV DNase/DNA nickases specific for phosphorothioated or glycosylated phage DNA. Within this conserved family is GmrSD, a Type IV modification dependent system which targets glucosylated 5hmC bases(28, 41). GmrS encodes a DUF262 domain while GmrD encodes a DUF1524 domain. DUF1524 contains a conserved HXXP motif that is seen in protein families such as the His-Me finger endonuclease superfamily(45). DUF262 is a member of the ParB-like superfamily that includes nucleases related to ParB in addition to uncharacterized proteins(41). Notably, the DUF262 region within GmrSD has been shown to encode a highly conserved DGQQR motif that is responsible for NTP binding and hydrolysis that could be important for DNA cleavage(41). This DGQQR motif is also found in both TgvA and TgvB, further supporting a functional link with GmrSD.

Studies have shown that a fused form of GmrSD in which the two domains are linked as one polypeptide is more abundant than systems where GmrS and GmrD are encoded as separate polypeptides(41). However, in either case, both GmrSD together are required for phage defense. TgvAB is distinct from this system in that TgvA only contains a DUF262 domain while TgvB contains both a DUF262 and DUF1524 region. Therefore, TgvB resembles a GmrSD fused polypeptide. Based on this, we predicted that TgvB alone should be sufficient for phage defense; however, expression of TgvA or TgvB alone with their native promoters or overexpressed using the pBAD promoter was insufficient for defense. Phage defense only occurred when both TgvA and TgvB were present. It is possible that TgvA and TgvB form a complex that is required for recognition and restriction of modified phage DNA. In support of this, the ORFs of *tgvA* and *tgvB* overlap by 8 basepairs, further suggesting a connection either through regulation of expression or some other mechanism. However, more studies need to be done to elucidate the molecular mechanism of TgvAB and how it specifically targets glucosylated 5hmC.

GmrSD targets and restricts glucosylated 5hmC phage DNA(28), and we report a similar result for TgvAB using both T2 and T4. Our data show that removing the *agt* (α-glucosyltransferase) and *bgt* (β-glucosyltransferase) genes responsible for glucosylating 5hmC in T4, T4 was able to bypass TgvAB defense (Fig. 5C). This protection can then be partially restored if the mutant T4 is passaged in a host overexpressing either T4 *agt* or *bgt* (Fig. 6C). Moreover, by evolving T2 in the presence of TgvAB, we identified 4 independent null mutations in the T2 *agt* gene that provide resistance to TgvAB protection. Interestingly, all these mutations occurred in poly-T tracts, suggesting these sites might be contingency loci allowing for rapid evolution of genome modification in response to different evolutionary pressures imposed by restriction enzymes, an idea which we are further exploring.

α- and β-glucosylation of 5hmC occurs in all T-even phage including T2, and *agt* and *bgt* are encoded by both T2 and T4. However, TgvAB resistant mutations in T2 were only observed in *agt,* suggesting that α-glucosylation is critical for T2 sensitivity to TgvAB. Our results indicating that the T4 ^Δ*agt*Δ*bgt*^ mutant passaged in a strain overexpressing T4*^bgt^* restored partial susceptibility clearly demonstrates that TgvAB can also target β-glucosylated 5hmC. Further studies are needed to deduce how TvgAB targets α- and/or β-glucose modification to 5hmC in different phage.

*E. coli* strain MG1655 encodes two known Type IV restriction systems, McrA and McrBC(46, 47). McrA and McrBC both separately target 5hmC and 5mC modifications, respectively(46, 47). We demonstrated that there is a trade-off for mutating *agt,* as losing the glucose modification to 5hmC renders T2 and T4 susceptible to recognition and restriction by other Type IV restriction systems such as McrA and McrBC. This suggests that phage can transition between glucosylation and unglucosylated DNA to evade different Type IV modification systems (Fig. 7).

Restriction modification (R-M) systems are one of the oldest known phage defense systems as observations of R-M systems have been made since the 1950s(23). Since then, multiple classes of R-M systems have been observed. These include Type I, Type II, and Type III systems that modify the host DNA and target unmodified phage DNA and the Type IV restriction enzymes that target unique modifications on phage DNA(23). Although all of these systems have been extensively studied, they have yet to be explored in depth in *V. cholerae* beyond a few limited computational studies that predicted R-M systems in *V. cholerae*(48, 49).

This study identifies a novel Type IV restriction system in *V. cholerae*. *V. cholerae* encodes three major lytic phages, ICP1, ICP2, and ICP3(3). However, none of these phages encode *agt* or *bgt*, and thus we predict they would be resistant to TgvAB. Therefore, resistance to TgvAB is one potential explanation for the rise to dominance of the ICP phage. However, it remains possible that TgvAB protects *V. cholerae* from infection by other undiscovered phages, and it would be interesting to search for phage that can infect a Δ*tgvAB V. cholerae* mutant by are restricted in the WT strain. Thus, additional studies to elucidate the function of TgvAB in its native context in *V. cholerae* and its impacts on phage infection are warranted.

## Methods

### Strains and growth conditions

All bacterial and phage strains are listed in Supplementary Table 1. Unless otherwise stated, cultures were grown in Lysogeny Broth (LB) at 37 °C in a shaking incubator set at 210rpm and supplemented with the following antibiotics for plasmid maintenance: tetracycline (10µg mL^-1^), kanamycin (100µg mL^-1^), chloramphenicol (100µg mL^-^1) and carbenicillin (100µg mL^-^1) as necessary. Ectopic expression was induced with arabinose (0.5mM) or IPTG (100µM). Coliphages were propagated by streaking on MMB agar (LB + 0.1mM MnCl^2^ + 5mM CaCl^2^ + 5mM MgCl^2^ + 0.5% agar) with *E. coli* DH10β at a 1:1,000 dilution of an overnight culture and incubated overnight. These plates were then covered with 10mL of phage buffer (0.1mM Tris HCl 7.5pH + 10mM MgSO^4^ + 0.4% NaCl^2^ + dH^2^O) and incubated statically at 4°C for 48 hours. Phage buffer was then filtered through a 0.22µM filter. Overnight cultures were prepared in the same media used in the experiment.

### Screening of a *Vibrio cholerae* cosmid library

The *V. cholerae* pLAFR cosmid library was previously described(18). Overnight cultures of the cosmid library in *E. coli* strain DH5⍺ were serially plated at dilutions of 10^-4^, 10^-5^, and 10^-6^ on LB + Tetracycline agar plates. Isolated colonies were grown in a 96-well plate with LB + Tetracycline at 37 °C in a shaking incubator overnight. Overnight cultures of the library and pLAFR(50, 51) (a broad host range, tetracycline-resistant cosmid cloning vector) were then mixed 1:1,000 with 5 mL of MMB agar + Tetracycline and poured into 4-well Nunc Rectangular Dishes (ThermoFisher). Dishes were dried at room temperature for 1 hour, then 5µl of 10-fold serial dilutions of T2 stored in LB were spotted, air dried and incubated overnight at 37 °C. Plaques from individual colonies were compared to plaques on pLAFR. All cosmids conferring T2 protection were extracted as described in plasmid construction and gene identification was confirmed using Oxford Nanopore Technology through the sequencing company, Plasmidsaurus.

### Efficiency of plaquing assay

Overnight cultures with appropriate plasmids were mixed 1:1,000 with 20mL of MMB agar + corresponding antibiotic and poured into 150×100mm petri dishes. Petri dishes were dried at room temperature for 1 hour. 5µl of a 10-fold dilution of corresponding coliphage in LB were spotted on top of the agar. Plaques were visualized using a Proteinsimple AlphaImager HP. All experiments were performed independently at least three times.

### Plaque forming unit quantification

Coliphage titer was determined using plaque assay as previously described(52). Overnight cultures with appropriate plasmids were mixed with the lowest concentration of coliphage (180µl of culture with 20µl of phage) where plaques were still visualized. Culture and phage were then mixed 1:1,000 with 20mL of MMB agar + corresponding antibiotic into 100×15mm petri dishes. Once plates solidified, they were incubated at 37°C overnight. Plaques were counted the next day. Data presented was calculated using Welch’s *t*-test and is representative of 6 biological replicates for pJBG007 and 3 biological replicates for p*tgvAB*. 3 technical replicates were done for each biological replicate.

### Growth curves assays

Overnight cultures with appropriate plasmids were diluted to an OD^600^ of 0.01 into LB supplemented with antibiotics. Cultures were grown for ∼3 hours to an OD^600^ of 0.1. Once cultures reach OD^600^ of 0.1, cultures were aliquoted into a 96-well plate and infected at an MOI of 0.01 in LB at 37 °C. Growth was monitored by measuring OD^600^ every 2.5 min for 8 hours using a BioTek 800 TS (Agilent) with continuous, linear shaking. Data presented are the mean and standard error of 4 biological replicates each with 3 technical replicates.

### Phage evolution experiments

Overnight cultures of p*tgvAB* and pMJF103 were diluted to an OD^600^ of 0.01 into LB + Kanamycin. Cultures were grown for ∼3 hours to an OD^600^ of 0.1 then split into 10 separate flasks, 5 containing p*tgvAB* and 5 containing pMJF103. Cultures infected with 10-fold serial dilutions in LB of T2 with the highest MOI at 10. Flasks were incubated for 4 hours at 37°C in a shaking incubator at 210 rpm. After 4 hours, the cultures were pooled and filtered using a 0.22µM filter. Overnight culture of p*tgvAB* was mixed 1:1,000 with 20mL MMB agar + Kanamycin and dried at room temperature for 1 hour. T2 lysates passaged on p*tgvAB* and pMJF103 were streaked for isolated plaques and placed at 37°C overnight. 5 isolated plaques from streak plates were picked, resuspended in 1mL of phage buffer and left static at 4°C for 24 hours. Overnight culture of p*tgvAB* was mixed 1:1,000 with 20mL MMB agar + Kanamycin and dried at 23°C for 1 hour. Isolated plaques suspended in phage buffer were streaked and placed at 37°C overnight. 10mL of phage buffer was added to each plate and left static at 4°C for 48 hours. 5µl of a 10-fold dilution of each T2 lysates was spotted on top of the MMB agar mixed 1:1,000 with p*tgvAB* to test for resistance. All lysates mentioned were passaged as mentioned above one more time on pMJF103. All lysates were used for further analysis and sequencing as described below. The culture passaged on only pMJF103 was also analyzed.

To extract phage DNA, 500μl phage lysates suspended in phage buffer were incubated at 98°C for 10 minutes to burst open the phage capsid head and inactivate any residual DNase and RNase from the bacterial culture. Samples were further purified using Wizard Genomic DNA Purification Kit (Promega) following the protocol for Isolation of Genomic DNA from Gram Positive and Gram-Negative Bacteria starting at step 3, under Lyse Cells. The resulting DNA was PCR-amplified for *agt* using oJBG0129 and oJBG0120 and *bgt* genes using oJBG0127 and oJBG0128. PCR products were cleaned and purified using the DNA Clean & Concentrator kit (Zymo Research) and Oxford Nanopore sequenced. Sequencing results were mapped to T2 reference genome (NC_054931.1) using Geneious Prime 2023.0.2.

### Plasmid construction

All primers are listed in Supplementary Table 3 and plasmids are listed in Supplementary Table 2. All PCR products were amplified using Q5-High Fidelity DNA polymerase according to the manufacturer (New England Biolabs).

pMJF103 was constructed as an arabinose-inducible vector that could be conjugated. pEVS141 vector backbone encoding the kanamycin resistance marker, p15A *oriV* and *oriT* was amplified with primers oMJF043 and oMJF044. An arabinose-inducible cassette including *araC* and mOrange under the pBAD promoter was amplified from the construct pBAD_mOrange (a gift of Patricia Champion) using Q5 and primers oMJF041 and oMJF042. Amplifications were confirmed by gel electrophoresis and treated with DpnI enzyme to remove template. Products were mixed 1:1 and transformed into chemically competent DH5α *E. coli* and plated on LB agar with 100 µg/mL kanamycin.

Transformants were picked for overnight broth culture in LB supplemented with 100 µg/mL kanamycin and 0.01% L-arabinose. Cultures showing mOrange expression were subsequently miniprepped using a Wizard miniprep kit (Promega). To remove the mOrange insert and generate and empty cloning vector, plasmid was double digested with NdeI and SpeI restriction enzymes (NEB) followed by gel purification of the 4725 bp band using a Wizard spin column (Promega). A multiple cloning site linker was prepared by annealing the primers oMJF055 and oMJF056 at 8 µM each in a buffer of 50 mM Tris HCl pH7.5, 50 mM NaCl and 1 mM EDTA. Linker was ligated into purified linearized vector using T4 ligase (NEB) at 4°C overnight followed by transformation into DH5α *E. coli*. Presence and orientation of the linker was confirmed by colony PCR using GoTaq polymerase (Promega) with primers oMJF042 and oMJF055. Colonies with confirmed insert were grown overnight in LB broth with 100 µg/mL kanamycin and plasmids were purified by miniprep.

To generate the p*tgvAB*, *tgvAB* was amplified from pJBG007 with 520bp upstream sequence to include its native promoter and overlapping ends to the pMJF103 using oJBG122 and oJBG106. To generate p*tgvA, tgvA* was amplified from pJBG007 with 520bp upstream sequence to include its native promoter and overlapping ends to pMJF103 using oJBG0122 and oJBG108. To generate *ptgvB, tgvB* was amplified from pJBG029 with 520bp upstream sequence to include its native promoter and overlapping ends to pMJF103 using oJBG122 and oJBG106. To generate T2 p*agt, agt* was amplified from T2 wild type with overlapping ends to pMJF103 using oJBG137 and oJBG138. To generate T4 p*agt, agt* was amplified from T4 wild type with overlapping ends to pMJF103 using oJBG133 and oJBG134. To generate T4 p*bgt, bgt* was amplified from T4 wild type with overlapping ends to pMJF103 using oJBG131 and oJBG132. The pMJF103 backbone was linearized using primers oMJF019 and oMJF053. All PCR products were cleaned and purified using the DNA Clean & Concentrator kit (Zymo Research).

All plasmid constructs were made using fast cloning(53). Amplified inserts and linearized pMJF103 backbone were DpnI treated for at 37°C for 1 hour followed by heat inactivation at 80°C for 20 min. Following DpnI treatment, 1µl of corresponding insert + 1µl of linearized pMJF103 backbone + 50µl of DH5α chemical competent cells were mixed and left on ice for 30 min. After 30 min. tubes were incubated at 42°C for 1 min. and immediately placed on ice for 3 min. 500µl of S.O.C was added, and cells were recovered at 37°C for 1 hour. Cells were spread on LB agar + Kanamycin and left at 37°C overnight. Inserts were verified by PCR amplification of the cloning sight using oMJF057 and oMJF058 primers.

To extract plasmids, Wizard *Plus* SV Minipreps DNA Purification System kit (Promega) was used following the Centrifugation Protocol under Quick Protocol. Miniprepped plasmids were introduced into DH10β cells by electroporation. All plasmids were confirmed by sequencing (Plasmidsaurus). Plasmid sequencing results were mapped to *V. cholerae* C6706 Chromosome I (CP064350) or T2 reference genome (NC_054931.1) using Geneious Prime 2023.0.2.

### Transposon mutant library

A transposon mutant library in pJBG007 was made using EZ-Tn5 <KAN-2> Insertion Kit (Lucigen). The following components were used for a 10µl reaction (EZ-Tn5 10X Reaction Buffer (1µl) + 1µg pJBG007 cosmid DNA (1µl) + EZ-Tn5 <KAN-2> Transposon undiluted (1µl) + EZ-Tn5 Transposase (1µl) + Nuclease Free H^2^O (6µl)). Reaction was incubated at 37°C for 2 hours. After 2 hours, 1µl of stop solution was add and reaction was incubated at 70°C for 10 min. 2µl of reaction was dialyzed for 15 min and then electroporated in 20µl of DH10β cells. Transformed cells were resuspended in 500µl of S.O.C (2% Tryptone + 0.5% yeast extract + 10mM NaCl + 2.5mM KCl + 10mM MgCl^2^ + 10mM MgSO^4^ + 20mM glucose) and recovered at 37°C shaking 210 rpm for 1 hour. 200µl of recovered culture and 5 individual 1:10 dilutions were plated on LB + Kanamycin + Tetracycline agar plates and incubated at 37°C overnight. All colonies were scraped and resuspended in 750µl of LB. 250µl of 80% glycerol was added to the 750µl resuspension and frozen at -80°C. Screening of Transposon Library was done as described in.

### Gene deletion in pJBG007

Lambda red recombineering was used as previously described(54–56). DH10β pJBG007 were made electrocompetent as follows: An overnight culture of pJBG007 grown in Yeast Extract and Nutrient Broth (YENB) (0.75% Bacto Yeast Extract + 0.80% Nutrient Broth, pH 7.5 using NaOH) + Tetracycline was back diluted 1:100 into YENB + Tetracycline and grown to an OD^600^ of ∼0.5. Cells were washed 3 times at 4°C with 10% glycerol. Cells were resuspended in 10% glycerol, aliquoted and stored at -80°C. pKD46(57) was dialyzed for 15 minutes and electroporated into pJBG007 electrocompetent cells. 500µl of S.O.C was added to cells and recovered at 30°C for 20 min. Cells were plated on LB agar + Tetracycline + Carbenicillin and incubated at 30°C overnight. pJBG007 + pKD46 cells were grown in LB + Tetracycline + Carbenicillin and made electrocompetent as described above. The chloramphenicol acetyltransferase (*cat*) cassette was PCR amplified using from pKD3(57) with overlapping ends to the either *tgvA* using oJBG037 and oJBG038, *tgvB* using oJBG022 and oJBG023, or *tgvAB* using oJBG047 and oJBG048. All PCR products were cleaned and purified using the DNA Clean & Concentrator kit (Zymo Research). The amplified chloramphenicol cassette was dialyzed as previously described and electroporated into pJBG007 + pKD46 electrocompetent cells. 500µl of S.O.C was added to cells and recovered at 37C for 1 hour. Cells were plated on LB agar + Tetracycline + Chloramphenicol and incubated at 37°C overnight. The insert was verified by PCR amplification using primers using oJBG021 and oJBG030 that amplified the outside of the insertion site. Overnight cultures of BW29427 with pTL17 supplemented with 300 µg/mL of diaminopimelic acid (DAP) was mixed with an overnight of pJBG007 + CAM cassette at a 1:1 ratio (25µl each) and spotted on an LB Agar + DAP plate. The spot was dried and incubated at 37C for 3 hours. The culture was struck on LB agar + Chloramphenicol + Kanamycin plates. Overnights of pJBG007 + CAM cassette + pTL17(58) were back diluted 1:100 into LB + Chloramphenicol + Kanamycin + IPTG and incubated at 37°C for 2 hours. 10-fold dilution of culture was made and dilutions from 10^3^-10^9^ were plated on LB agar + Tetracycline plates at 37°C overnight.

Patch plating starting with LB agar + Tetracycline, LB agar + Chloramphenicol, LB agar + Kanamycin, and ending with LB agar, was used to confirm removal of CAM cassette and pTL17 from host. Deletions are left with an FRT scar after removal of CAM cassette. Deletions were confirmed by sequencing (Plasmidsaurus).

### Complementation of Phage AGT and BGT

Overnight cultures of T2 p*agt* and p*bgt* and T4 p*agt* and p*bgt* were diluted 1:1000 in LB + Kanamycin + Arabinose and incubated for ∼3 hours at 37°C. Cultures were then mixed 1:1,000 in 20mL MMB agar + Kanamycin + Arabinose and dried at room temperature for 1 hour. All T2 mutants (Mutant 2, 3, 4, 5, and 6) were streaked on both T2 p*agt* and p*bgt* and T4 ^Δ*agt*Δ*bgt*^ were streaked on both T4 p*agt* and p*bgt.* Plates were then incubated at 37°C. Plates were covered with 10mL phage buffer and left static at 4°C for 48 hours. Phage buffer was filtered through a 0.22µM filter. 5µl of a 10-fold dilution of each T2 mutants and T4 Δ*agt*Δ*bgt* that were passaged on their corresponding p*agt* or p*bgt* was spotted on top of the MMB agar mixed 1:1,000 with either p*tgvAB*, pMJF103, MG1655, or MG1655 Δ*mcrBC*Δ*mcrA*.

## ACKNOWLEDGEMENTS

This research was supported and funded by NIH grants GM139537 and AI158433 to C.M.W. We thank Mike Laub and Sriram Srikant for construction and sharing of the T4^D*agt*D*bgt*^ and MG1655 D*mcrA*D*mcrBC* mutants, Douglas Guzior for writing R script used for statistical analysis, and Micah Ferrell for plasmid construction of pMJF103 and subsequent primer construction. We thank the laboratory of Melanie Blokesch who independently discovered TvgAB for coordinating joint submission of this research.

## FIGURE LEGENDS

**Extended Data Fig. 1.**
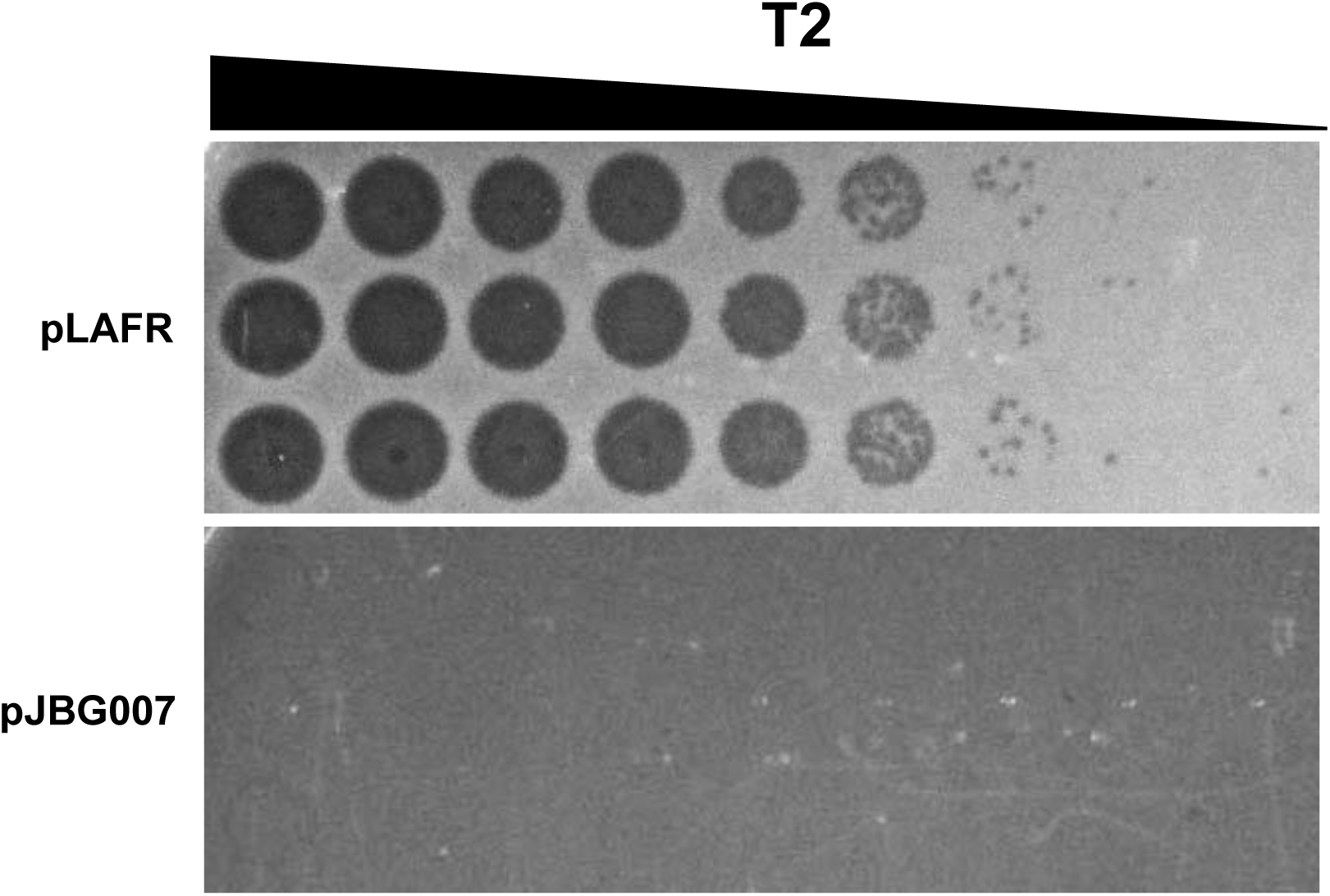
Efficiency of plaquing (EOP) of T2. 10-fold serial dilutions of T2 spotted on empty vector (pLAFR) and pJBG007 done in triplicate corresponding for Fig. 1A.

**Extended Data Fig. 2.**
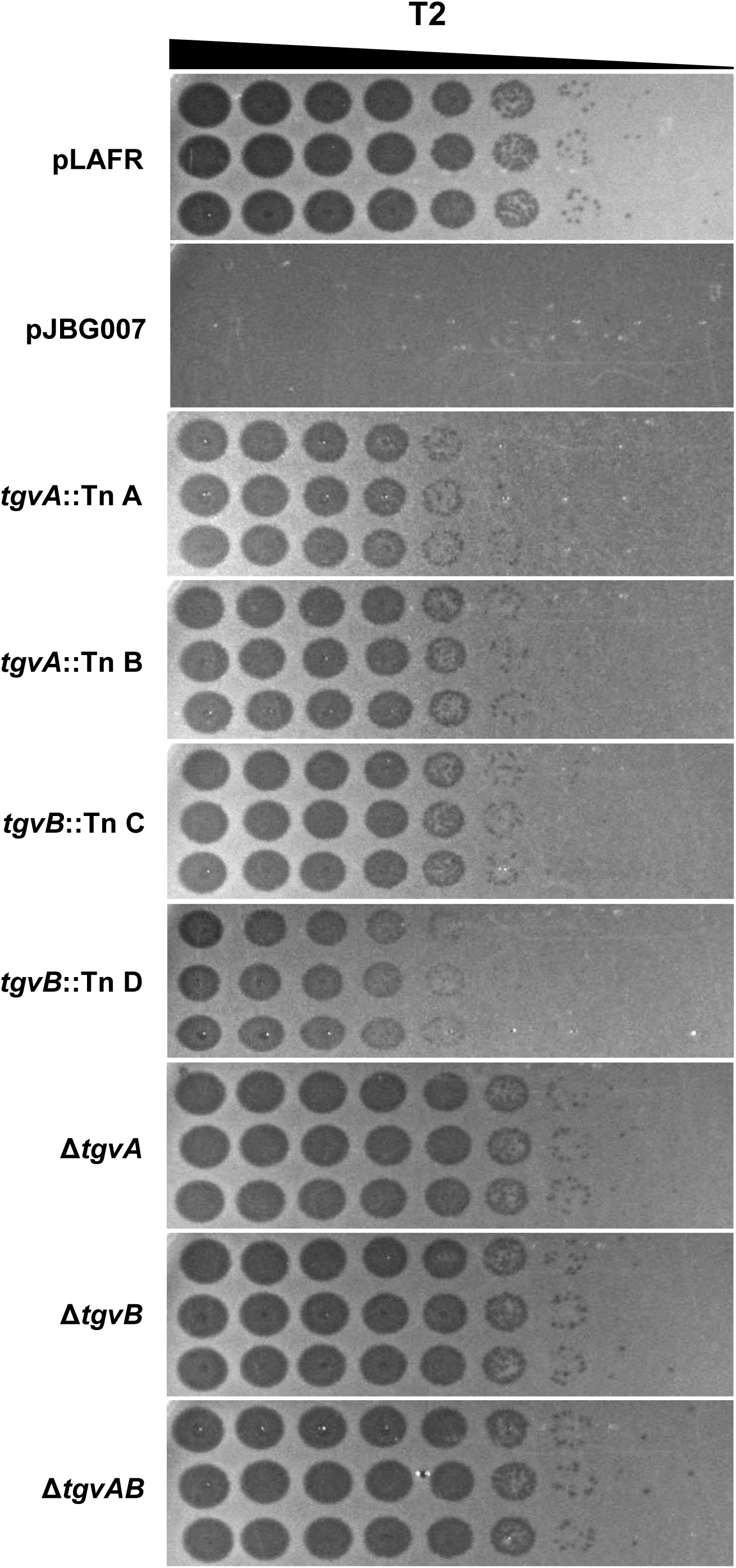
EOP of T2 on Tn and deletion host. 10-fold serial dilutions of T2 spotted on empty vector (pLAFR), pJBG007, p*tgvA*::Tn A, p*tgvA*::Tn B, p*tgvB*::Tn C, p*tgvB*::Tn D, Δ*tgvA,* Δ*tgvB,* and Δ*tgvAB* deletions on pJBG007 done in triplicate corresponding for Fig. 2A.

**Extended Data Fig. 3.**
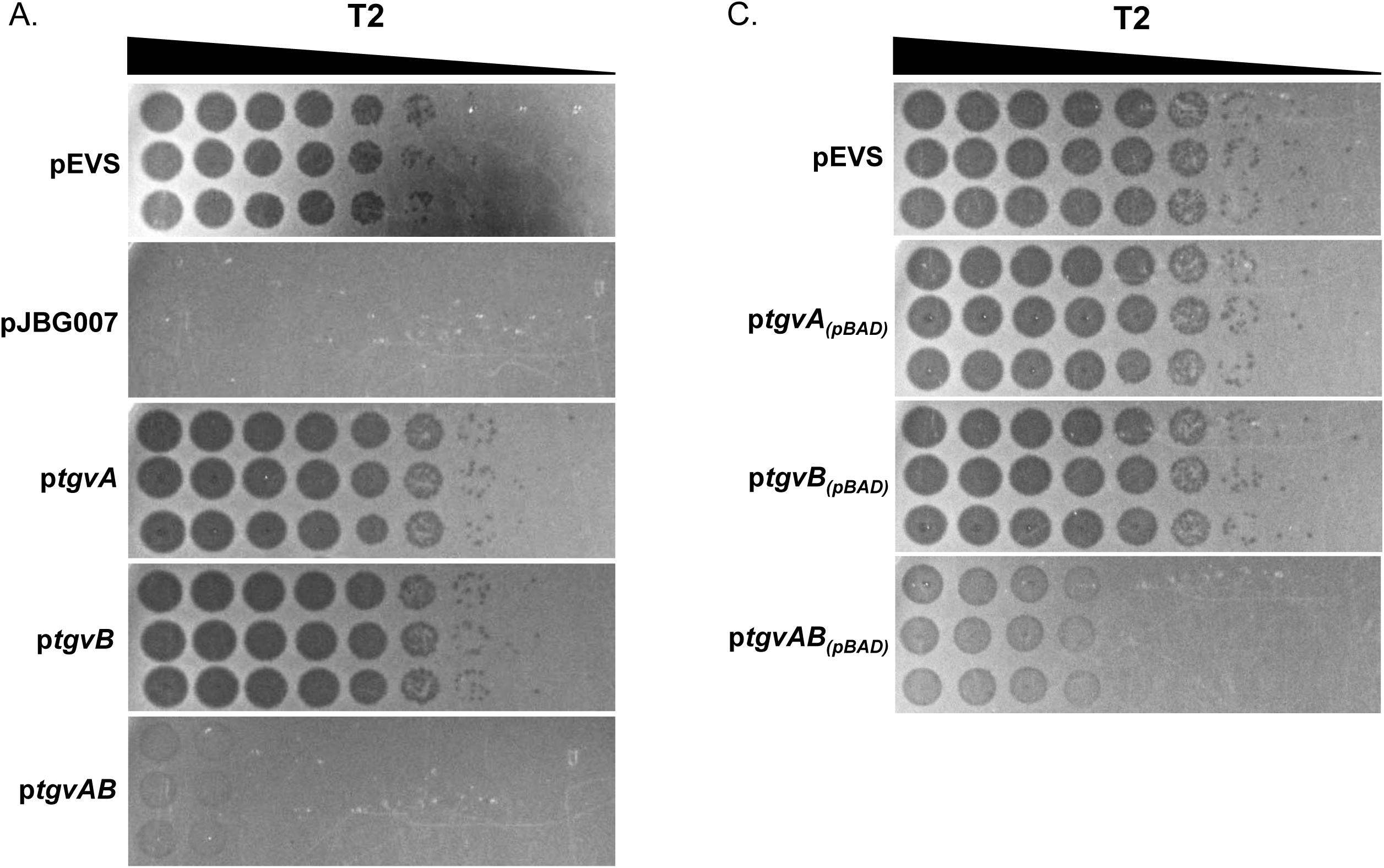
EOP of T2 on p*tgvA*, p*tgvB*, p*tgvAB*. **A.** 10-fold serial dilutions of T2 spotted on p*tgvA*, p*tgvB,* and p*tgvAB* with native expression done in triplicate corresponding to Fig. 3A. and **B.** p*tgvA****_(pBAD)_***, p*tgvB****_(pBAD)_***, and p*tgvAB****_(pBAD)_*** expressed using an arabinose inducible pBAD promotor done in triplicate corresponding to Fig. 3B.

**Extended Data Fig. 4.**
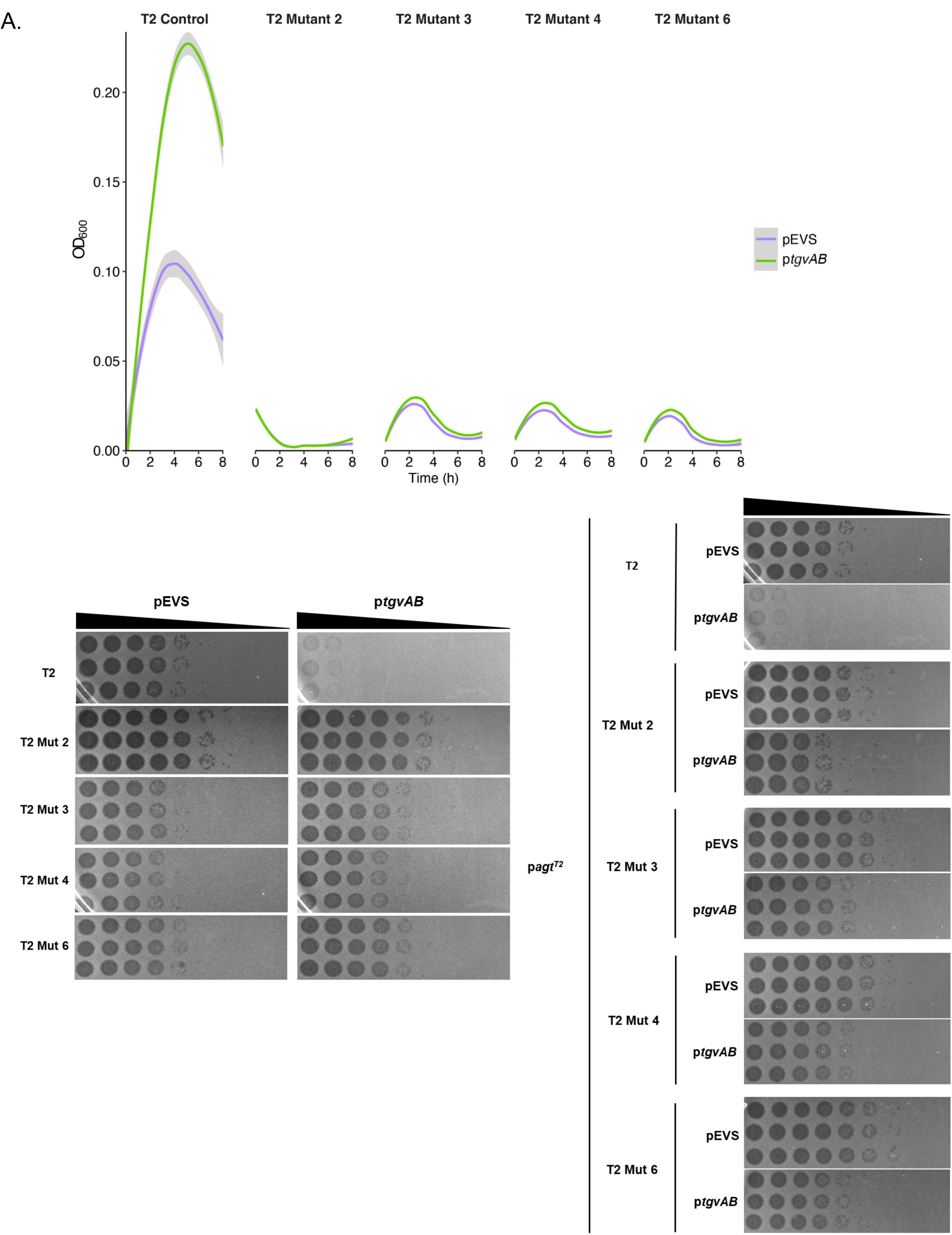
T2 mutants escape TgvAB defense. **A.** Growth curve of pEVS (empty vector) (purple line) and p*tgvAB* (green line) infected at time 0 with T2 Control, and all 4 T2 evolved mutants at an MOI of 0.01 is shown. The mean and standard error of 4 biological replicates each with 4 technical replicates are presented. **B.** 10-fold serial dilutions of WT T2 and all 4 evolved T2 mutants spotted on empty vector and p*tgvAB* done in triplicate corresponding to Fig. 4A. **C.** 10-fold serial dilutions of WT T2 and all 4 evolved T2 mutants passaged on host expressing p*agt*^T2^ with a pBAD promotor spotted on empty vector and p*tgvAB* done in triplicate corresponding to Fig. 4D.

**Extended Data Fig. 5.**
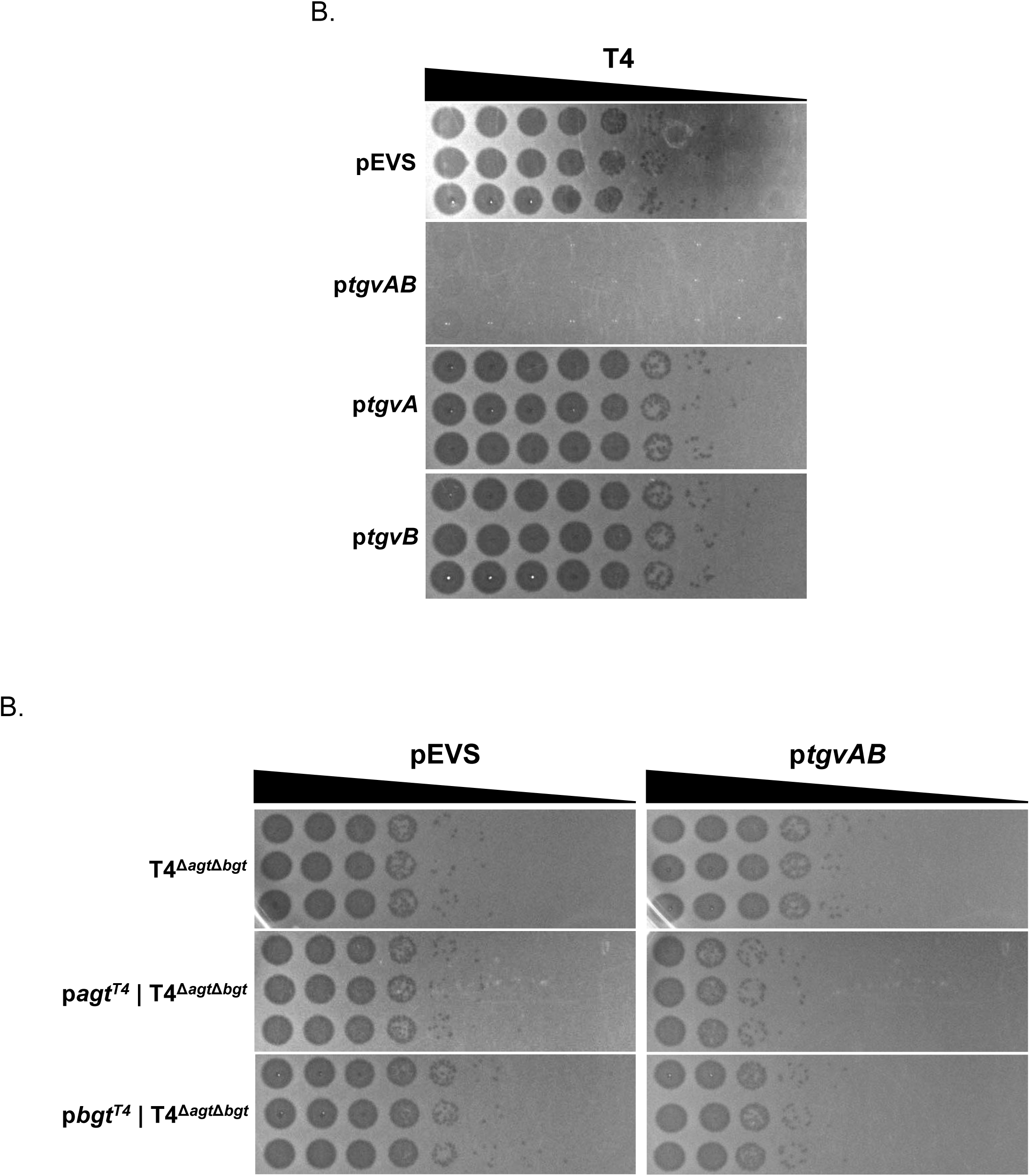
EOP of T4 and T4^Δ*agt*Δ*bgt*^. **A.** 10-fold serial dilutions of T4 spotted on empty vector, p*tgvA,* p*tgvB*, and p*tgvAB* done in triplicate corresponding with Fig. 5B. **B.** 10-fold serial dilution of T4*^ΔagtΔbgt^* (top), T4^Δ*agt*Δ*bgt*^ passed on host expressing p*agt*^T4^ (middle), or p*bgt*^T4^ (bottom) with a pBAD promotor spotted on empty vector and p*tgvAB* host done in triplicate corresponding to Fig. 5C.

**Extended Data Fig. 6.**
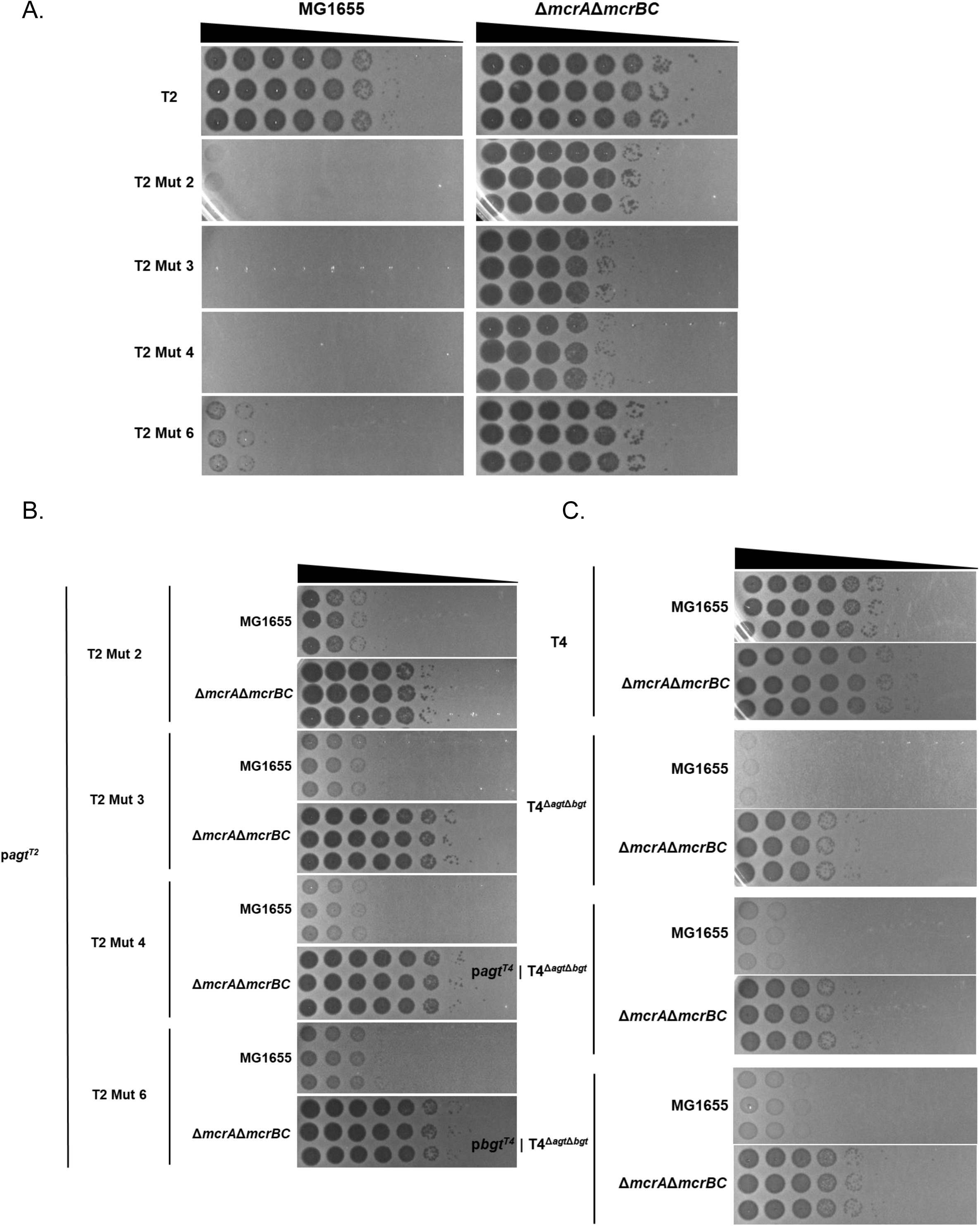
EOP of T2 Mutants on MG1655 and MG1655 Δ*mcrA*Δ*mcrBC*. **A.** 10-fold serial dilutions of WT T2 and all 4 T2 mutants spotted on WT MG1655 and MG1655 Δ*mcrA*Δ*mcrBC* done in triplicate corresponding to Fig. 6A. **B.** 10-fold serial dilutions of WT T2 and all 4 T2 mutants passaged on host expressing p*agt*^T2^ with a pBAD promotor spotted on WT MG1655 and MG1655 Δ*mcrA*Δ*mcrBC* done in triplicate corresponding to Fig. 6B. **C.** 10-fold serial dilution of WT T4, T4^Δ*agt*Δ*bgt*^, T4^Δ*agt*Δ*bgt*^ passed on host expressing p*agt*^T4^, or p*bgt*^T4^ with a pBAD promotor on WT MG1655 and MG1655 Δ*mcrA*Δ*mcrBC* host done in triplicate corresponding to Fig. 6C.

